# Conservation of endangered galaxiid fishes in the Falkland Islands requires urgent action on invasive brown trout

**DOI:** 10.1101/2021.06.15.448501

**Authors:** J. F. Minett, D.M. Fowler, J.A.H. Jones, P. Brickle, G.T. Crossin, S. Consuegra, C. Garcia de Leaniz

## Abstract

Non-native salmonids are protected in the Southern hemisphere where they sustain aquaculture and valuable sport fisheries, but also impact on native galaxiid fishes, which poses a conservation conundrum. Legal protection and human-assisted secondary releases may have helped salmonids to spread, but this has seldom been tested. We reconstructed the introduction of brown trout (*Salmo trutta*) to the Falkland Islands using historical records and modelled its dispersal. Our results indicate that establishment success was ∼88%, and that dispersal was facilitated by proximity to introduction sites and density of stream-road crossings, suggesting it was human assisted. Brown trout has already invaded 54% of Falkland rivers, which are 2.9-4.5 times less likely to contain native galaxiids than uninvaded streams. Without strong containment we predict brown trout will invade nearly all suitable freshwater habitats in the Falklands within the next ∼70 years, which might put native freshwater fishes at a high risk of extinction.

## Introduction

Invasive species represent one of the major threats to freshwater biodiversity, and yet their introduction has in many cases been intentional. For example, salmonids have been deliberated translocated all over the world to provide fishing and aquaculture opportunities since the 19^th^ century (McDowall 2006), despite being responsible for the demise of native fish fauna (Garcia de Leaniz et al. 2010; Young et al. 2010).

Human activities have not only been responsible for the introduction of invasive species, but have also helped in many cases with their expansion (Hulme 2015). Yet, the importance of human assisted dispersal of non-native species is often difficult to assess due to lack of accurate introduction records and confounding environmental factors (Tabak et al. 2017). Islands provide ideal scenarios to examine the dispersal of invasive species because the date and location of introductions are typically well known, and there is often baseline information on the status of native species before the invasion (Ewel and Högberg 1995).

Brown trout (*Salmo trutta*) is one of the most successful freshwater invaders and has been included as one of the ‘100 of the world’s worst invasive alien species’ (Lowe et al. 2000) due to its widespread ecological damage. The species has been implicated in the decline of native galaxiid fishes in many parts of the Southern hemisphere (McDowall 2006), most notably in South America (Elgueta et al. 2013; Young et al. 2010), New Zealand (McDowall 2003), and the Falkland Islands, where it has benefitted from protected status (McDowall et al. 2001). This has created a conservation conundrum because protecting non-native salmonids to boost sport fishing may have put native fish at risk (Garcia de Leaniz et al. 2010).

Three surveys, conducted 10 and 20 years ago, concluded that brown trout had severely impacted two of the three native galaxiids, *Aplochiton zebra* and *Aplochiton taeniatus* (Fowler 2013; McDowall et al. 2001; Ross 2009), which appear to have contracted their range and are threatened by secondary releases. However, little is known about the current distribution of the endangered galaxiids, or the roles that natural and human-mediated dispersal may have played in the dispersal of brown trout following the initial introductions.

Here we reconstructed the introduction and establishment of brown trout in the Falkland Islands using historical records and modelled its dispersal using anthropogenic and bioclimatic variables. We also derived risk maps under different management scenarios to help conservation agencies identify galaxiid populations at high risk of invasion and prioritise the establishment of freshwater refugia.

## Methods

### Reconstructing the introductions of brown trout

Historical records of the introduction of salmonids in the Falkland Islands (Table S1) were compiled and cross-referenced from (Arrowsmith and Pentelow 1965) and (Stewart 1973; Stewart 1980), supplemented with information from the media in the United Kingdom (Salmon and Trout 2012), and from (Basulto 2003), (Faundez et al. 1997) and (Daciuk 1975) in South America. We also searched the Stirling University library for shipment records from the Howietoun hatchery (built in 1881) as this had dispatched salmonid eggs across the world.

### Sampling of freshwater fish

A database of presence/absence records of the four species of freshwater fish present in the Falklands (three native galaxiids, *A. zebra, A. taeniatus*, and *G. maculatus;* and the non-native brown trout) was compiled by cross-checking records from (McDowall et al. 2001; Ross 2009), expanded with information from angler reports and our own sampling (Fowler 2013). McDowall’s (McDowall et al. 2001) fish survey employed seine, gill and fyke netting, spotlighting at night, and electrofishing of ∼50 m stream reaches. Ross (Ross 2009) employed electrofishing, seine netting and visual checks. We used single-pass electrofishing (Smith-Root ELBP2), seine netting and visual surveys during 2011-2013 to add 28 new sites to the database, bringing the total number of sites sampled for freshwater fish to 134 (Table S2).

### Species distribution modelling

We divided the Falkland Islands into 8,813 1×1 km^2^ grid cells, and excluded cells that had more than 30% of their area in the sea and those that contained no rivers, as in a previous species distribution modelling (Rodriguez-Rey et al. 2019). Brown trout presence was modelled using 13 anthropogenic and 9 bioclimatic predictors (Table S3) for which we extracted mean values or took the value from the centre of the grid cell using QGIS 3.4. We excluded three predictors with VIF>3 to reduce collinearity, and randomly divided equal numbers of presence and absence records into training and testing datasets from 134 sites (Table S2) to train and test the model.

Brown trout distribution was modelled via a generalized linear model and the Leave One Out Cross Validation (LOOCV) (Hooten and Hobbs 2015). The *drop1* function in R3.5.3 (R Core Team 2019) was used to test the significance of individual predictors and arrive at a minimal adequate model based on changes in AIC. Model performance was assessed using the under the curve (AUC) criterion and compared against a null model built using the same testing and training datasets used for the final model (Rodriguez-Rey et al. 2019) using parametric bootstrapping (1,000 simulations).

### Establishment success and calculation of invasion risk

To calculate establishment success, we compared the proportion of introduction sites that still had brown trout ∼50 years later against the random 50% expectation using a binomial test. We then used presence/absence data for brown trout and the three native galaxiids to assess how the presence of brown trout influenced the presence of native galaxiids by calculating relative risks. To visualize invasion risk, we used QGIS 3.10.3 to generate invasion risk maps based on the probability of brown trout occurrence, calculated using the LOOCV procedure across all valid grid cells, as explained above.

### Predictive modelling of brown trout invasions under different management scenarios

We modelled the occupancy of brown trout since 1950 and predicted its future dispersal over a 130-year period (∼30 generations) considering three different management scenarios: (1) no containment, (2) moderate containment (a 10% reduction in the probability of invasion at each cell), and (3) strong containment (a 30% reduction in the probability of invasion at each cell). For cells with a high probability of invasion (p ≥ 0.8) we assumed that containment would not be effective so we maintained the original invasion probabilities (i.e. no containment, scenario 1).

As grid cells were found to be more likely to become invaded if they were close to invaded sites (see Results), we calculated the Euclidean distance from each uninvaded site to the nearest invaded site. Invasion probabilities were then updated at each iteration under the three scenarios outlined above. Each scenario was run over 300 iterations to obtain a mean percentage occupancy and 95% binomial confidence intervals. We used the observed rate of expansion (0.9% increase in occupancy/yr since 1950) to calibrate the model and convert the number of model iterations into calendar years (one iteration = ∼24 years or ∼4 generations).

### Ethics & Permits

Fish sampling was carried out under permit number R18/2018 (17/04/2018) issued by the Falkland Islands Government (Falkland Islands Environmental Committee) and Swansea University Ethics Committee (Reference number SU-Ethics-Student-081217/307; SU-Ethics-Student-090118/299; SU-Ethics-Student-160118/463.

## Results

### Origin of brown trout

Approximately 113,000 brown trout eggs were shipped to the Falkland Islands on eight separate occasions over an 18-year period (1944-1962, Table S1). Although original records are missing, many consignments were described as arriving in ‘excellent condition’ (Stewart 1973). The first introductions (1944-1947) came from the Lautaro hatchery in Chile (Arrowsmith and Pentelow 1965; MacCrimmon and Marshall 1968), and were primarily sourced from resident (i.e. non-anadromous) parents of German origin (Faundez et al. 1997; Radcliffe 1922). Subsequent eggs came from three sources in England: Surrey, Pentlands and Lancashire. The Surrey and Pentlands fish were from resident parents, while the Lancashire trout were derived from ‘sea run trout’ caught in the River Lune (Arrowsmith and Pentelow 1965). The provenance of the Pentlands resident trout is unclear, but it may have originated from Cobbinshaw Loch (Arrowsmith and Pentelow 1965; Stewart 1973), Loch Leven (Fish Loch Leven 2019), or the Howietoun Hatchery (Ross Gardiner, pers. comm.). The Howietoun hatchery had reared trout from Loch Leven and many other sources, but we found no records of fish having ever been sent to the Falkland Islands. In total 28 different sites were stocked (Table S1), but three rivers – all within a 25 km radius of the capital Stanley - received most of the introductions.

### Establishment success

Of the 17 stocked sites for which there are fish survey data, 15 sites still had brown trout ∼50 years later. Establishment success can therefore be estimated as 88% (95CI = 62-98%), which is significantly better than chance (χ^2^ = 8.47, df = 1, *P* = 0.004).

### Modelling of brown trout distribution

At the time of the last survey (2012), brown trout occupied 54% of all sampled 1km^2^ grid cells, with *Aplochiton* spp. only occupied 18%, confined to the South of the Islands (Fig. 1). Brown trout occurrence was predicted by distance to the nearest invaded point in a straight line (estimate = -0.238, SE = 0.067, *t* = -3.56, *P* < 0.001), absence of *Aplochiton* spp. (estimate = -1.57, SE = 0.769, *t* = -2.04, *P* = 0.041) and number of river road crossings in the drainage basin (estimate = 0.156, SE = 0.066, *t* = 2.37, *P* = 0.018). The model explained the occurrence of brown trout significantly better than chance (LRT = 52.17, df = 3, *P* <0.001, AUC = 0.85).

**Figure 1.**
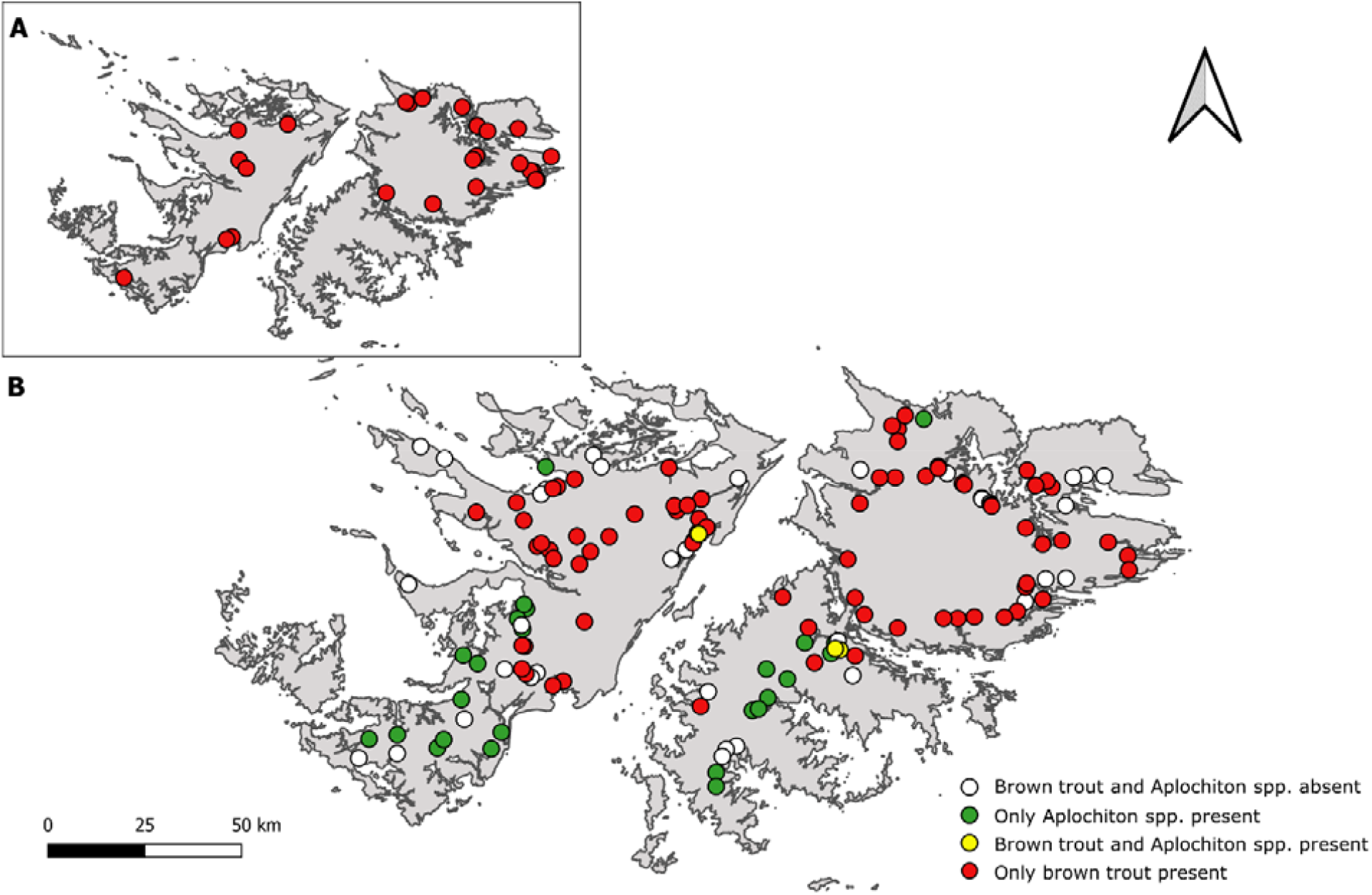
Map of the Falkland Islands showing (A) sites of the historical introductions of brown trout during 1944-1962 (details given in Table S1) and (B) presence/absence of brown trout and native *Aplochiton* sp. based on 1999-2012 surveys (detailed in Table S2) with 6 additional sites sampled in 2018-2019.

### Impact of brown trout on native galaxiids

Native galaxiids were less likely to occur in streams invaded by brown trout than in uninvaded ones (Fig. 2), but the impact of invasive brown trout was more pronounced in the case of *Aplochiton sp*. Calculation of relative risk indicated that *Aplochiton sp* was 4.5 times less likely to persist in streams invaded by brown trout than in uninvaded streams (95CI = 1.8-11.2, *P*<0.001). For *Galaxias maculatus*, the presence of brown trout decreased the probability of occurrence 2.9 fold (95CI = 2.0-4.2, *P*<0.001).

**Figure 2.**
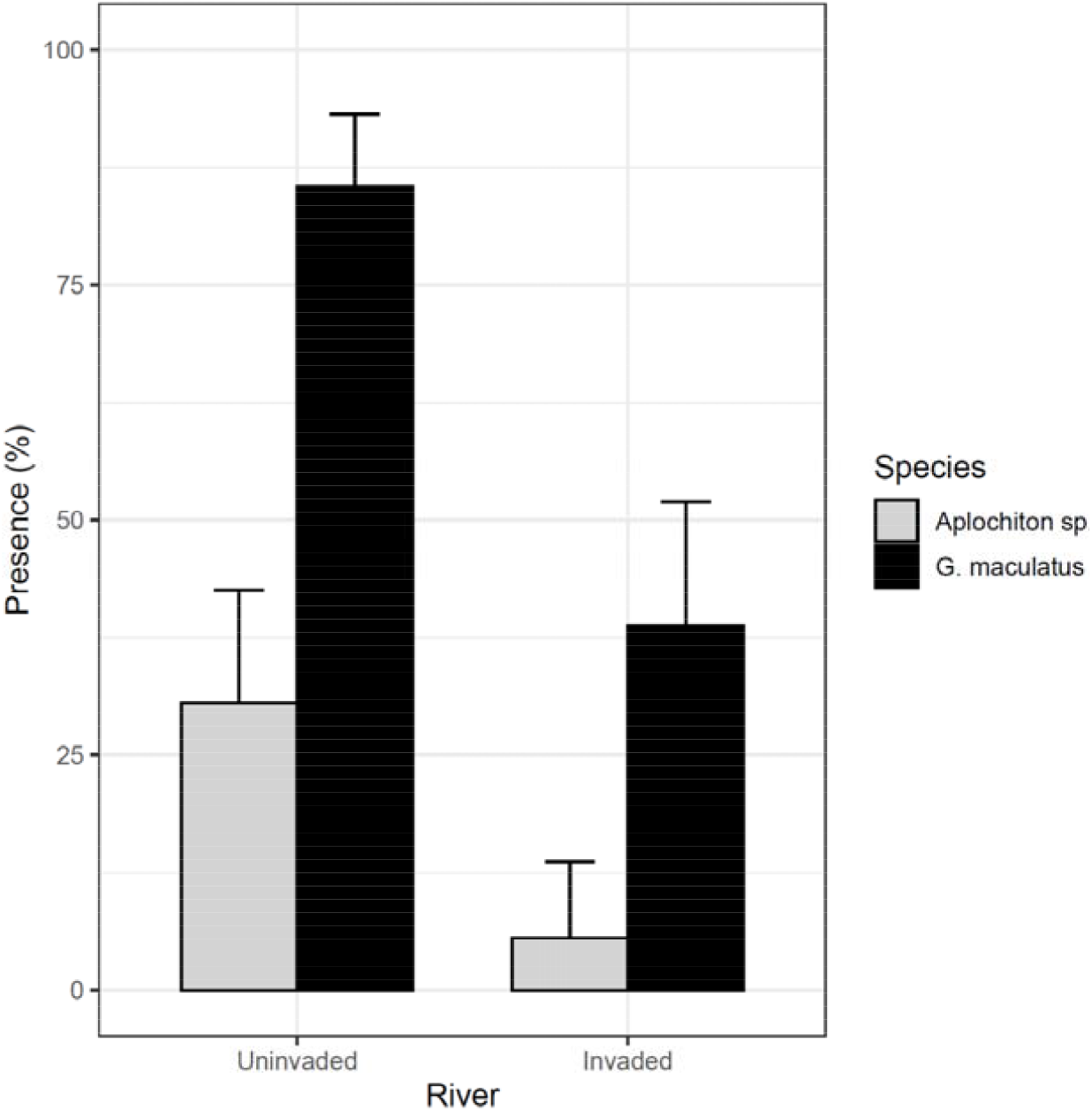
Frequency of occurrence (% and binomial upper 95CI) of native galaxiids (*Galaxias maculatus* and *Aplochiton* sp.) in streams invaded by brown trout (N=62) and in uninvaded streams (N= 72).

### Invasion risk

A risk map generated from the probability of occurrence of brown trout identified 21% of cells with a very high risk of invasion (R≥0.75), 24% with a high risk (R = 0.50-0.75), 17% with moderate risk (R=0.25-0.50) and 38% with low risk (R<0.25). By overlaying the distribution of the endangered *Aplochiton* spp. we identified 9 sites at a high or very high risk of invasion where preventive measures should be prioritised to exclude brown trout and protect native freshwater fish (Fig. 3).

**Figure 3.**
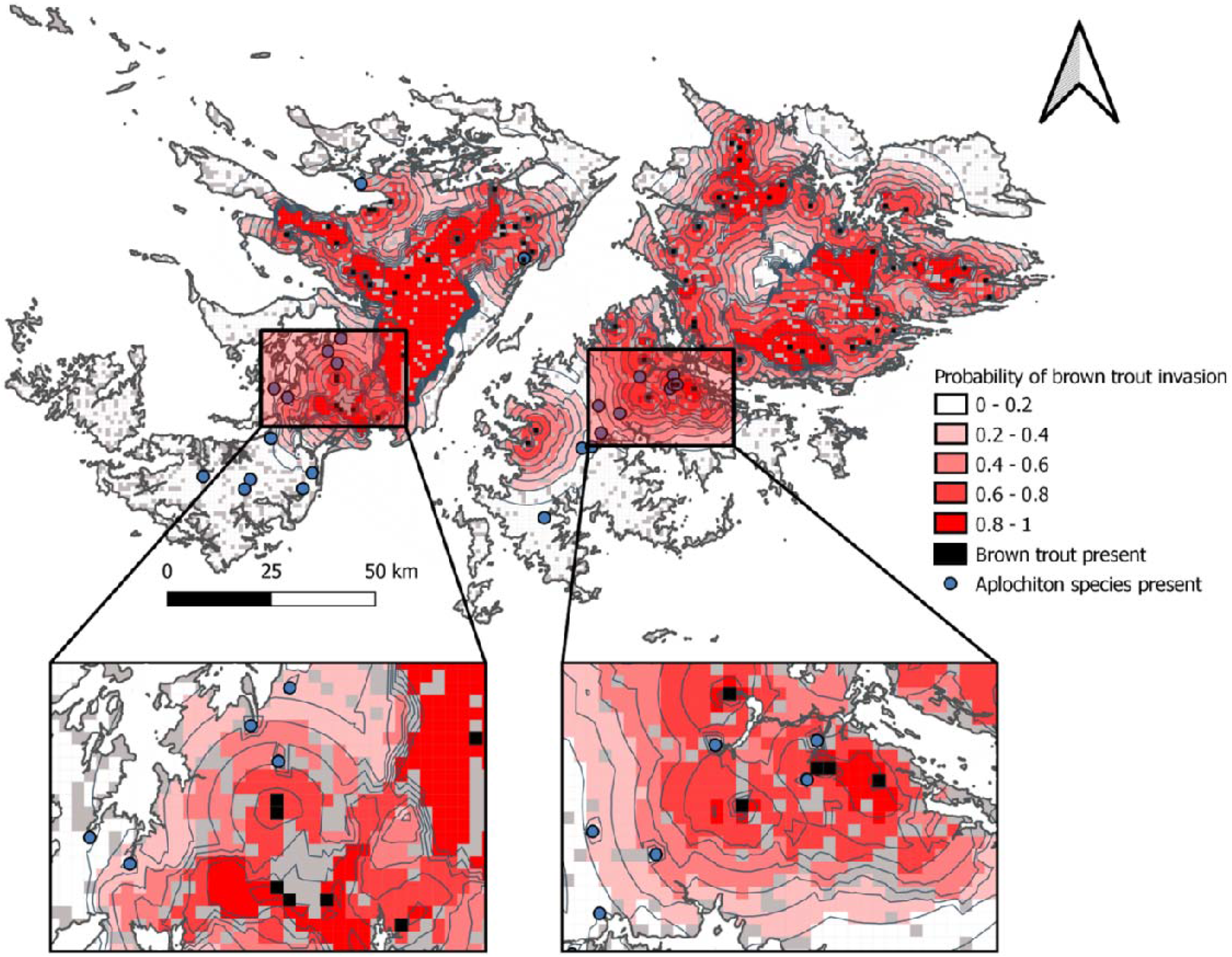
Risk maps showing probabilities of brown trout invasion based on species distribution modelling. *Aplochiton* refugia at a high risk of brown trout invasion are shown in the zoomed insets.

### Invasion under different management scenarios

Our simulations indicate that if nothing is done (scenario 1) brown trout will likely increase its occupancy from 54% to 93% within the next 70 years (95CI = 70-99%). Under scenario 2 (moderate containment) occupancy is predicted to increase to 86% (95CI = 59-94%) and to 69% (95CI = 47-81%) with strong containment (scenario 3, Fig. 4). Thus, occupancy is predicted to increase under all the three scenarios, but only with strong containment can current *Aplochiton* refugia be protected from trout invasions.

**Figure 4.**
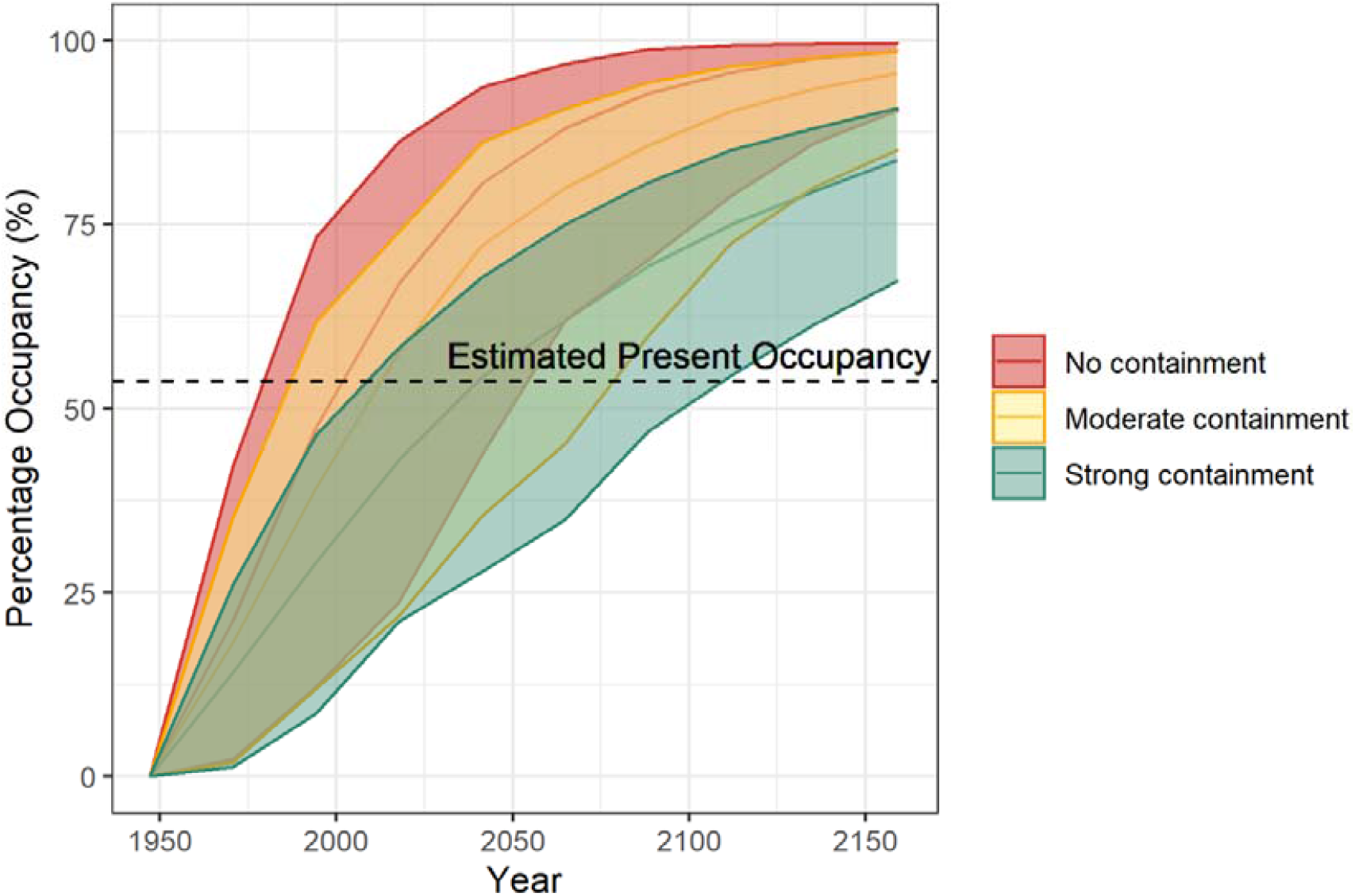
Modelled expansion of brown trout in the Falkland Islands under three different management scenarios.

## Discussion

Our study indicates that brown trout have already invaded 54% of the streams in the Falkland Islands since they were introduced in 1944-1962, and are impacting on native freshwater fish. Streams invaded by brown trout were 2.9 times less likely to harbour *Galaxias maculatus* and 4.5 times less likely to contain *Aplochiton* sp. than uninvaded streams, suggesting the impacts are substantial. This finding is consistent with competitive exclusion of native galaxiids by invasive brown trout (Garcia de Leaniz et al. 2010; Young et al. 2009), exacerbated by predation and trophic interference (Arismendi et al. 2014; Elgueta et al. 2013). Our simulations suggest that unless more stringent measures are put in place, brown trout will likely invade nearly all the suitable freshwater habitats in the Falklands within the next ∼70 years. Given that endangered *Aplochiton* sp. only occupy ∼18% of the area, mostly confined to southern part of the Islands, this could drive the species to extinction.

The establishment success of brown trout in the Falklands was very high (88%), as seen elsewhere in the Southern Hemisphere (Arismendi et al. 2014; Young et al. 2010). For example, in Argentina no failed introduction of brown trout was ever reported (Baigún and Quirós 1985). Several factors may help explain this. Firstly, our study shows that brown trout introduced into the Falklands originated from at least four different origins with two life history strategies, which resulted in genetic admixture (Minett et al. 2021b). Multiple origins and genetic admixture can increase genetic diversity and facilitate adaptation to novel conditions (Consuegra et al. 2011), which along with repeated introductions may increase invasion success. Another reason for the success of brown trout in the Falklands may lie in their high phenotypic plasticity and facility for marine dispersal, which seems to have facilitated the colonization of coastal rivers in the Falklands, a point recently confirmed by acoustic tracking (Minett et al. 2021b).

However, marine dispersal alone cannot explain the current distribution of brown trout in the Islands; secondary translocations must have also taken place because the species is now found in land-locked sites, where it could not have reached without human intervention. Transporting brown trout has been illegal in the Falklands since 1999, but some translocations must have taken place (McDowall et al. 2001). Indeed, our results indicate that brown trout presence was predicted by proximity to other invaded sites (overland, but not around the coast) and by the density of river-road crossings, which is consistent with secondary translocations facilitated by the road network, as seen in many other aquatic invasive species. For example, roads facilitated the expansion of smallmouth bass (*Micropterus dolomieu*) in remote lakes in Canada (Kaufman et al. 2009) and of bluegill (*Lepomis macrochirus*) in Japan (Kizuka et al. 2014). The Falklands has ∼800 km of road tracks crisscrossing a dense river network, most of which have been built over the last three decades (Fowler and Garcia de Leaniz 2012), and it is likely that this may have facilitated the expansion of brown trout. Recent eDNA analysis of water samples (Minett et al. 2021a) has revealed the presence of brown trout in additional streams since our last survey, suggesting that the species is expanding at a rate of ∼0.9%/year.

Other invasive salmonids are also threatening the native fish fauna of the Falklands. For example, both chinook salmon (*Oncorhynchus tshawytscha*) and coho salmon (*Oncorhynchus kisutch*) are increasingly being caught off West Falkland (Fowler 2013), most likely originating from Chile or Argentina, highlighting the potential for further salmonid invasions. Similarly, the development of sea trout farming in open-net cages in the Falklands in 2013 poses a risk of escapees, which could further compromise the survival of native galaxiids, as seen in Patagonia (Consuegra et al. 2011; Vanhaecke et al. 2012), particularly if sea cages are located close to *Aplochiton* refugia. Given the widespread ecological damage caused by invasive salmonids, being able to identify areas at high risk of invasion is critical for managing and curtailing their expansion. In this sense, our risk maps allows conservation officers to identify high risk areas, and could be used as part of an integrated management strategy for managing invasive salmonids in the Falklands.

### Conclusions & Recommendations

Galaxiids rank among the most severely threatened fish in the world due to the introduction of invasive salmonids (Garcia de Leaniz et al. 2010; McDowall 2006). Our modelling suggests that without containment and strict measures brown trout will likely invade all remaining suitable water bodies in the Falklands before the end of the century, putting the endangered freshwater fish of the Islands at a high risk of extinction.

Existing legislation makes it illegal to transport or propagate brown trout in the Falklands, but this seems insufficient as the species is also afforded a protected status, and fishing for trout is widely promoted (Falkland Islands Government 2015), which may facilitate its spread. The road network appears to be a main route of human-assisted translocations, and is essential that more stringent measures are put in place. This may involve making people more aware of the impacts of salmonid invasions and passing more stringent legislation. Exclusion barriers could also be deployed around galaxiid refugia to reduce the risk of salmonid invasions (Jones et al. 2021b), but care must be taken to ensure this does not impact on native galaxiids, which may pose a challenge as even small barriers can have negative impacts on weak swimmers (Jones et al. 2021a). Brown trout is subject to a bag limit and a strict fishing season and it might be useful to consider lifting these restrictions in some places to slow down the invasion front. Intensive fishing could be used to eradicate brown trout and establish buffer zones around *Aplochiton* refugia; analysis of eDNA from water samples could be used to delineate such refugia (Minett et al. 2021a), to serve as an early warning of brown trout invasions, and to establish whether containment measures have been successful.

Since McDowall’s call for action 20 years ago (McDowall et al. 2001), brown trout has continued to expand while native galaxiids have continued to decline. *Aplochiton* sp. features in a postal stamp of the Islands while *Galaxias maculatus* is called ‘Falklands minnow’, testifying to their importance for local islanders, and their place in the natural and cultural heritage of the Islands. Salmonids have brought wealth and recreation opportunities to the Falklands, but have also caused the demise of native freshwater fish. Our study indicates that urgent protection measures are needed to safeguard their survival.

## Acknowledgements

We thank William P. Kay for help calculating distance around the coast, Supercomputing Wales for use of the Sunbird cluster, and Falkland Islands Government IMS-GIS Centre and SAERI for providing GIS data. Funding from Fortuna Ltd. and Swansea University College of Science is gratefully acknowledged.

---

**Table S1.**
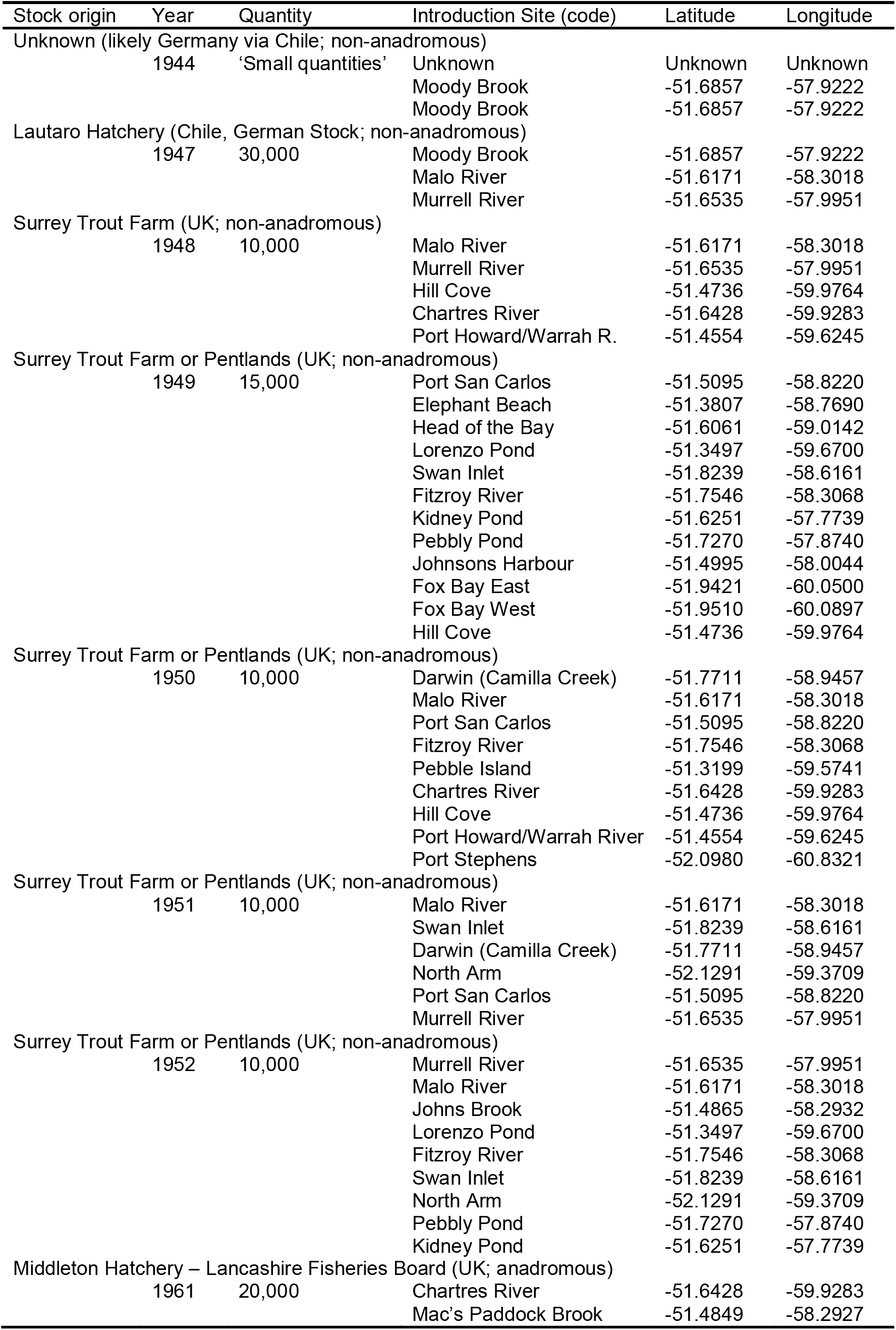
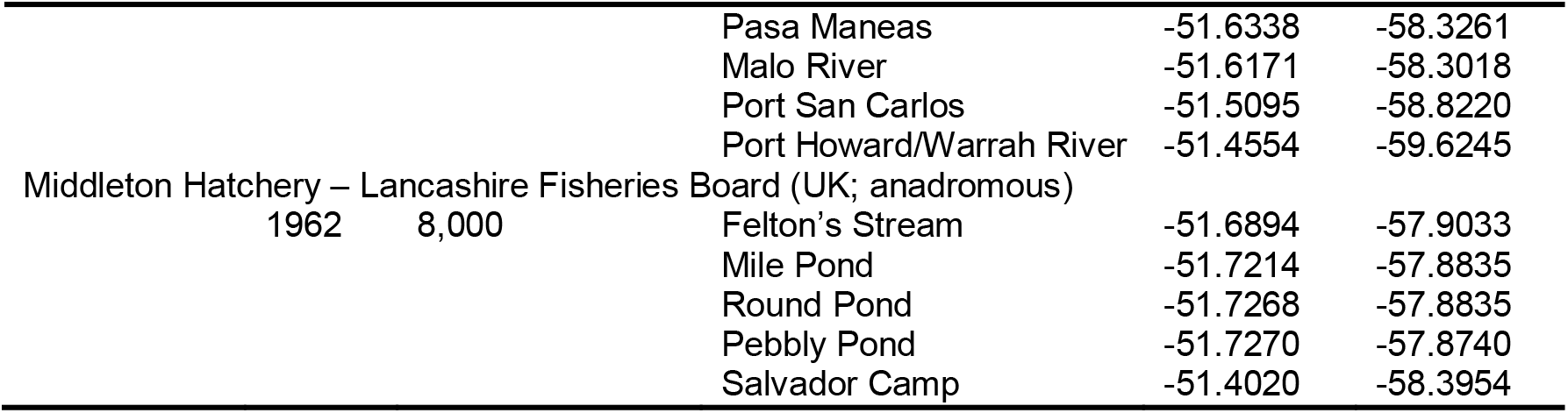
Introduction of brown trout eggs in the Falkland Islands, 1944-1962.

**Table S2.** Presence/absence of brown trout (*St*), *Aplochiton* sp. (*Ap*) and *Galaxias maculatus* (*Gm*) in the Falkland Islands. Sites marked with an asterisk denote brown trout introduction sites (see Table S1).

| Sample Site | Island | Date last sampled | Lat. | Long. | *St* | *Ap* | *Gm* | Reference |
| --- | --- | --- | --- | --- | --- | --- | --- | --- |
| Arrow Harbour Arroyo | East | 2011 | -51.9062 | -58.9508 | + | - | + | Fowler (2013) |
| Arroyo Pedro | East | 1999 | -51.5111 | -58.5346 | + | - | - | McDowall et al. (2001) |
| Big Pond | East | 2018 | -51.8433 | -51.7522 | + | - | + | Minett (unpubl.) |
| Bodie Creek | East | 2011 | -51.9191 | -59.1042 | + | - | + | Fowler (2013) |
| Bull Pass Stream | East | 2011 | -51.8909 | -59.0074 | + | + | + | Fowler (2013) |
| Clay Ditch | East | 1999 | -51.4925 | -58.6700 | + | - | - | McDowall et al. (2001) |
| Colorado Pond | East | 2018 | -51.7136 | -58.4717 | + | - | - | Minett (unpubl.) |
| Comoda Ditch | East | 1999 | -51.8219 | -58.5003 | + | - | - | McDowall et al. (2001) |
| Congo Ponds Area | East | 2009 | -51.9787 | -59.5072 | - | - | - | Ross (2009) |
| Dan's Shanty Stream | East | 1999 | -51.5236 | -58.2020 | + | - | + | McDowall et al. (2001) |
| Darwin (Camilla Creek) * | East | 1999 | -51.7711 | -58.9457 | + | - | - | McDowall et al. (2001) |
| Deep Arroyo | East | 2011 | -51.9553 | -59.2080 | - | + | + | Fowler (2013) |
| Ditches into New Haven | East | 2009 | -51.7646 | -59.2166 | + | - | + | Ross (2009) |
| Duffins Bridge Stream | East | 2009 | -52.1078 | -59.4073 | - | - | + | Ross (2009) |
| Elephant Beach Pond* | East | 1999 | -51.3807 | -58.7690 | + | - | + | McDowall et al. (2001) |
| Elmer's Ditch Tributary | East | 1999 | -51.4939 | -58.7840 | + | - | - | McDowall et al. (2001) |
| Estancia Brook | East | 2008 | -51.6480 | -58.1678 | + | - | - | Fowler (2013) |
| Felton Stream* | East | NA | -51.6894 | -57.9033 | NA | NA | NA | Not sampled |
| Findley Creek Stream | East | 2011 | -51.8882 | -59.0250 | + | + | + | Fowler (2013) |
| Findlay Harbour (Wreck House) | East | 2009 | -52.0122 | -59.5375 | + | - | - | Fowler (2013) |
| Fitzroy River* | East | 2012 | -51.7546 | -58.3068 | + | NA | NA | Angler |
| Frying Pan | East | 2011 | -51.8111 | -58.3387 | + | - | - | Fowler (2013) |
| Gonzales Arroyo | East | 2009 | -51.9513 | -58.9621 | - | - | + | Ross (2009) |
| Green Pass Brook Tributary | East | 1999 | -51.4098 | -58.7715 | + | - | - | McDowall et al. (2001) |
| Head of the Bay* | East | 2012 | -51.6061 | -59.9764 | + | NA | NA | Angler |
| Head of the Creek Stream | East | 1999 | -51.4961 | -58.0763 | - | - | + | McDowall et al. (2001) |
| Hunter's Arroyo | East | 1999 | -52.1147 | -59.4470 | - | - | + | McDowall et al. (2001) |

**Table S2 (cont).**
| Sample Site | Island | Date last sampled | Lat. | Long. | St | Ap | Gm | Reference |
| --- | --- | --- | --- | --- | --- | --- | --- | --- |
| John's Brook* | East | 2018 | -51.4834 | -58.2930 | + | - | NA | Minett (unpubl.) |
| Johnsons Harbour* | East | NA | -51.4995 | -58.0044 | NA | NA | NA | Not sampled |
| Kidney Pond* | East | NA | -51.6251 | -57.7739 | NA | NA | NA | Not sampled |
| Laguna Isla | East | 2011 | -51.8439 | -58.7890 | + | - | - | Fowler (2013) |
| L'Antioja Stream | East | 2012 | -51.8249 | -58.5627 | + | - | - | Fowler (2013) |
| Little Creek Stream | East | 1999 | -51.3504 | -58.7418 | + | - | - | McDowall et al. (2001) |
| Lorenzo Pond* | East | 1973 | -51.3593 | -58.6730 | - | + | + | Stewart (1973) |
| Mac's Paddock Brook* | East | NA | -51.4849 | -58.2927 | NA | NA | NA | Not sampled |
| Magellan Pond | East | 1999 | -51.4995 | -58.0062 | - | - | + | McDowall et al. (2001) |
| Malo River* | East | 1999 | -51.6171 | -58.3018 | + | - | + | McDowall et al. (2001) |
| Mary Hill Quarry | East | 2018 | -51.6844 | -57.7894 | + | - | - | Minett (unpubl.) |
| Mile Pond* | East | NA | -51.7214 | -57.8835 | NA | NA | NA | Not sampled |
| Moody Brook* | East | 2012 | -51.6857 | -57.9222 | + | - | - | Fowler (2013) |
| Mullet Creek | East | 1999 | -51.7187 | -57.9185 | + | - | + | McDowall et al. (2001) |
| Murrell River* | East | 2013 | -51.6535 | -57.9951 | + | - | - | Fowler (2013) |
| North Arm* | East | 2011 | -52.1291 | -29.3709 | - | - | + | Fowler (2013) |
| Northwest Arm House Stream | East | 2018 | -52.1674 | -59.4874 | - | + | + | Minett (unpubl.) |
| Northern Stream | East | 1999 | -51.5018 | -58.1223 | - | - | + | McDowall et al. (2001) |
| Orequta Arroyo | East | 2011 | -51.8373 | -59.1229 | + | - | - | Fowler (2013) |
| Pasa Maneas* | East | NA | -51.6338 | -58.3261 | NA | NA | NA | Not sampled |
| Pebbly Pond* | East | NA | -51.7270 | -57.8740 | NA | NA | NA | Not sampled |
| Round Pond* | East | NA | -51.7268 | -57.8835 | NA | NA | NA | Not sampled |
| Rumford Brook | East | 2011 | -51.6557 | -58.2399 | + | - | - | Fowler (2013) |
| Salvador Camp* | East | NA | -51.4020 | -58.3954 | NA | NA | NA | Not sampled |
| San Carlos River at Ford* | East | 2018 | -51.5095 | -58.220 | + | - | - | Minett (unpubl.) |
| Shepherds Brook | East | 1999 | -51.6808 | -58.9688 | + | - | + | McDowall et al. (2001) |
| Spots Arroyo | East | 2018 | -52.0260 | -59.3432 | - | + | + | Minett (unpubl.) |

| Stream at Colorado Pass | East | 2009 | -51.8702 | -59.0137 | - | - | + | Ross (2009) |
| --- | --- | --- | --- | --- | --- | --- | --- | --- |
| **Table S2 (cont).** |  |  |  |  |  |  |  |  |
| Sample Site | Island | Date last sampled | Lat. | Long. | St | Ap | Gm | Reference |
| Stream at Gibraltar Gate | East | 1999 | -51.4933 | -58.8417 | + | - | - | McDowall et al. (2001) |
| Stream near Hunter Arroyo | East | 1999 | -52.1303 | -59.4622 | - | - | + | McDowall et al. (2001) |
| Stream, Caneja Creek | East | 1999 | -51.5190 | -58.2621 | + | - | - | McDowall et al. (2001) |
| Stream, at Fitzroy | East | 1999 | -51.7838 | -58.2425 | + | - | + | McDowall et al. (2001) |
| Stream, Douglas Creek | East | 1999 | -51.4704 | -58.6229 | + | - | - | McDowall et al. (2001) |
| Stream, Elephant Beach Pond* | East | 2018 | -51.3731 | -58.7911 | + | - | + | Minett (unpubl.) |
| Stream, Fitzroy | East | 1999 | -51.7887 | -58.3085 | - | - | - | McDowall et al. (2001) |
| Stream, Monty Dean's Creek | East | 1999 | -51.5669 | -58.1515 | - | - | + | McDowall et al. (2001) |
| Stream, Mount Pleasant | East | 1999 | -51.8243 | -58.3878 | + | - | - | McDowall et al. (2001) |
| Stream, NW of Teal Inlet | East | 1999 | -51.5456 | -58.4638 | - | - | + | McDowall et al. (2001) |
| Stream NW of Teal Inlet | East | 1999 | -51.5481 | -58.4629 | - | - | + | McDowall et al. (2001) |
| Stream, Salt House Creek | East | 2009 | -52.1997 | -59.4894 | - | + | + | Ross (2009) |
| Stream, SE of Teal Inlet | East | 1999 | -51.5580 | -58.4330 | - | - | - | McDowall et al. (2001) |
| Stream, Smylie's Brook | East | 1999 | -51.4738 | -58.9129 | - | - | - | McDowall et al. (2001) |
| Stream, Teal Inlet | East | 1999 | -51.5654 | -58.4278 | + | - | - | McDowall et al. (2001) |
| Stream, SE of Teal Inlet | East | 1999 | -51.5622 | -58.4294 | - | - | + | McDowall et al. (2001) |
| Swan Inlet* | East | 2012 | -51.8239 | -58.6161 | + | NA | NA | Angler |
| Teal Creek Arroyo | East | 2009 | -51.8106 | -58.9121 | + | - | + | Ross (2009) |
| Third Corral Brook | East | 1999 | -51.5520 | -58.9182 | + | - | - | McDowall et al. (2001)) |
| Trib., Halfway House Arroyo | East | 2011 | -51.9977 | -59.2836 | - | + | + | Fowler (2013) |
| Turners Stream | East | 1999 | -51.5137 | -58.5277 | + | - | - | McDowall et al. (2001) |
| Unnamed Stream | East | 1999 | -51.8726 | -59.1386 | - | + | + | McDowall et al. (2001) |
| Unnamed Stream | East | 1999 | -51.9311 | -59.2855 | - | + | + | McDowall et al. (2001) |
| Unnamed Stream | East | 1999 | -51.4869 | -58.5905 | - | - | - | McDowall et al. (2001) |
| Unnamed Stream | East | 1999 | -52.0228 | -59.3197 | - | + | + | McDowall et al. (2001) |
| Unnamed Stream | East | 1999 | -51.8987 | -59.0414 | - | + | + | McDowall et al. (2001) |
| Unnamed Stream | East | 1999 | -51.8753 | -59.0254 | - | + | + | McDowall et al. (2001) |

| Unnamed Stream | East | 1999 | -51.7461 | -58.3020 | + | - | - | McDowall et al. (2001) |
| --- | --- | --- | --- | --- | --- | --- | --- | --- |
| **Table S2 (cont).** |  |  |  |  |  |  |  |  |
| Sample Site | Island | Date last sampled | Lat. | Long. | St | Ap | Gm | Reference |
| Unnamed Stream | East | 1999 | -51.7366 | -58.2306 | - | - | + | McDowall et al. (2001) |
| Unnamed Stream | East | 1999 | -51.7357 | -58.1542 | - | - | - | McDowall et al. (2001) |
| Unnamed Stream | East | 1999 | -51.4747 | -58.6230 | + | - | - | McDowall et al. (2001) |
| Pebble Island* | Pebble | NA | -51.3199 | -59.5741 | NA | NA | NA | Not sampled |
| 1<sup>st</sup> Arroyo | West | 2009 | -52.0838 | -60.5346 | - | + | - | Ross (2009) |
| 2<sup>nd</sup> Pass Stream | West | 1999 | -51.6253 | -60.1260 | + | - | - | McDowall et al. (2001) |
| 2<sup>nd</sup> Arroyo | West | 2009 | -52.0647 | -60.5092 | - | + | + | Ross (2009) |
| Arroyo Chico | West | 1999 | -51.9208 | -60.1891 | + | - | - | McDowall et al. (2001) |
| Arroyo Malo | West | 1999 | -51.9194 | -60.1471 | - | - | - | McDowall et al. (2001) |
| Ballan Stream | West | 2009 | -51.6471 | -59.5714 | - | - | + | Ross (2009) |
| Beach Stream | West | 1999 | -51.6258 | -59.5402 | + | - | + | McDowall et al. (2001) |
| Bull Hill Stream | West | 1999 | -51.5764 | -59.5194 | + | - | - | McDowall et al. (2001) |
| Bull Stream | West | 1999 | -51.4888 | -60.0413 | + | - | - | McDowall et al. (2001) |
| Campbell Creek Stream | West | 2009 | -52.0197 | -60.4278 | - | - | + | Ross (2009) |
| Cemetery Creek Estuary | West | 1999 | -51.5961 | -59.4913 | + | - | + | McDowall et al. (2001) |
| Chartres River* | West | 1999 | -51.6428 | -59.9283 | + | - | - | McDowall et al. (2001) |
| Daddy's Ditch | West | 2009 | -51.4195 | -59.9033 | - | - | + | Ross (2009) |
| Dean's River | West | 2009 | -52.0914 | -60.6869 | - | - | + | Ross (2009) |
| Dirty Ditch at the High Tide | West | 1999 | -51.5401 | -60.3470 | + | - | + | McDowall et al. (2001) |
| Double Stream | West | 2009 | -51.6677 | -59.6277 | - | - | + | Ross (2009) |
| Doyle River Tributary | West | 1999 | -51.7695 | -60.1764 | - | + | + | McDowall et al. (2001) |
| Dunbar Creek | West | 2009 | -51.4124 | -60.4556 | - | - | + | Ross (2009) |
| Edye Creek | West | 2012 | -51.8711 | -60.4206 | - | + | + | Fowler (2013) |
| Fish Creek 1 | West | 2012 | -52.0538 | -60.2908 | - | + | + | Fowler (2013) |
| Fish Creek 2 | West | 2018 | -51.8918 | -60.3681 | - | + | + | Minett (unpubl.) |
| Fox Bay East* (Doctors Creek) | West | 2018 | -51.9421 | -60.0500 | + | - | NA | Minett (unpubl.) |
| Fox Bay West* | West | 2012 | -51.9510 | -60.0897 | + | NA | NA | Angler |

| Gibraltar Stream<br>**Table S2 (cont).** | West | 2009 | -52.0913 | -60.3318 | - | + | + | Ross (2009) |
| --- | --- | --- | --- | --- | --- | --- | --- | --- |
| Sample Site | Island | Date last sampled | Lat. | Long. | St | Ap | Gm | Reference |
| Green Hill Stream | West | 2009 | -51.5552 | -59.6012 | + | - | - | Ross (2009) |
| Herbert Stream | West | 1999 | -51.5222 | -60.1959 | + | - | - | McDowall et al. (2001) |
| Hawk's Nest Stream | West | 1999 | -51.8055 | -59.9614 | + | - | - | McDowall et al. (2001) |
| Hill Cove* | West | 2008 | -51.4736 | -59.9764 | + | - | - | Fowler (2013) |
| House Creek | West | 1999 | -51.6097 | -59.5267 | + | + | + | McDowall et al. (2001) |
| House Stream | West | 2012 | -51.6116 | -59.5221 | + | + | + | Fowler (2013) |
| Lake Sullivan, Outflow | West | 2009 | -51.7922 | -60.2111 | - | + | + | Ross (2009) |
| Lake Sullivan North | West | 2011 | -51.8167 | -60.1941 | - | + | + | Fowler (2013) |
| Lake Sullivan North Outflow | West | 1999 | -51.8073 | -60.1976 | - | - | + | McDowall et al. (2001) |
| Lake Sullivan South | West | 2011 | -51.8567 | -60.1891 | + | - | + | Fowler (2013) |
| Larger Stream, Hill Cove | West | 1999 | -51.4945 | -60.0843 | - | - | + | McDowall et al. (2001) |
| Leicester Stream | West | 2009 | -51.9084 | -60.2682 | - | - | + | Ross (2009) |
| Main Point Creek | West | 2009 | -51.4474 | -59.8757 | - | - | + | Ross (2009) |
| Many Branches Stream | West | 1999 | -51.5303 | -59.5083 | + | - | + | McDowall et al. (2001) |
| Mickey Doolan's Ditch | West | 1999 | -51.9292 | -60.1703 | - | - | + | McDowall et al. (2001) |
| Mt Adam Stream | West | 1999 | -51.6355 | -60.0802 | + | - | - | McDowall et al. (2001) |
| Mt Donald Pond Inflow | West | 1999 | -51.5643 | -60.1720 | + | - | - | McDowall et al. (2001) |
| Neil Clark Nature Reserve | West | 2018 | -51.6324 | -59.5452 | + | - | NA | Minett (unpubl.) |
| Poncho Valley Stream | West | 2012 | -51.9736 | -60.4357 | - | + | + | Fowler (2013) |
| Port Howard/Warrah* | West | 1999 | -51.4554 | -59.6245 | + | - | - | McDowall et al. (2001) |
| Port Stephens* | West | 2009 | -52.0980 | -60.8321 | - | - | + | Ross (2009) |
| River Doyle | West | 2009 | -51.7588 | -60.1845 | - | + | + | Ross (2009) |
| Rous Creek Stream | West | 2009 | -51.7006 | -60.6122 | - | - | + | Ross (2009) |
| Skull Pass Stream | West | 1999 | -51.5444 | -59.6095 | + | - | - | McDowall et al. (2001) |
| Stewart's Brook | West | 2009 | -52.0482 | -60.6826 | - | + | + | Ross (2009) |
| Stream by Mt Rosalie House | West | 2009 | -51.4856 | -59.3685 | - | - | + | Ross (2009) |
| Stud Paddock Stream | West | 1999 | -51.9416 | -60.0497 | - | - | + | McDowall et al. (2001) |

| Teal House River | West | 2018 | -51.6194 | -60.1103 | + | - | NA | Minett (unpubl.) |
| --- | --- | --- | --- | --- | --- | --- | --- | --- |
| **Table S2 (cont).** |  |  |  |  |  |  |  |  |
| Sample Site | Island | Date last sampled | Lat. | Long. | *St* | *Ap* | *Gm* | Reference |
| Teal Ponds & Waterfall Stream | West | 1999 | -51.6557 | -60.0672 | + | - | + | McDowall et al. (2001) |
| Top Hog Ground Stream | West | 1999 | -51.6711 | -59.9714 | + | - | - | McDowall et al. (2001) |
| Unnamed Stream | West | 1999 | -51.5443 | -59.5602 | + | - | - | McDowall et al. (2001) |
| Unnamed Stream | West | 1999 | -51.9092 | -60.2030 | + | - | - | McDowall et al. (2001) |
| Unnamed Stream | West | 2012 | -51.6098 | -59.8563 | + | - | - | Fowler (2013) |
| Unnamed Stream, Hill Cove | West | 1999 | -51.5053 | -60.1034 | - | - | + | McDowall et al. (2001) |
| Unnamed Stream, Hill Cove | West | 1999 | -51.4928 | -60.0587 | + | - | - | McDowall et al. (2001) |
| Warrah River | West | 1999 | -51.5598 | -59.7581 | + | - | - | McDowall et al. (2001) |
| Waterfall Stream | West | 2009 | -51.3817 | -60.5421 | - | - | + | Ross (2009) |
| Weedy Outlet | West | 1999 | -51.8548 | -60.1980 | + | - | + | McDowall et al. (2001) |
| West Lagoons | West | 2013 | -51.4421 | -60.0817 | - | + | + | Fowler (2013) |
| Whisky Creek | West | 2009 | -52.0546 | -60.7896 | - | + | - | Ross (2009) |

**Table S3.** Predictors used to generate a species distribution model of brown trout in the Falkland Islands. Predictors shown in **bold** had a VIF< 3 and were included in the model.

| **Predictor** | **Description** | **Source** |
| --- | --- | --- |
| Anthropogenic predictors |  |  |
| **Eucl_dist_inv** | Euclidean distance to nearest invaded site in a straight line (km) | Own creation |
| **Coast_dist_inv** | Distance to nearest invaded site around the coast (km) | Own creation |
| **Eucl_dist_intro** | Euclidean distance to nearest introduction site in a straight line (km) | Own creation |
| **Coast_dist_intro** | Distance to nearest introduction site around the coast (km) | Own creation |
| **Intro_site** | Introduction site (y/n) | Table S1 |
| Intro_basin | Introduction basin (y/n) | Table S1 |
| **Settle_dist** | Distance to nearest settlement (km) | Own creation, FIG IMS-GIS Centre |
| **Road_dist** | Distance to nearest road (km) | Own creation, FIG IMS-GIS Centre |
| **Road_cross_no** | Number of river-road crossings in the river basin | Own creation, FIG IMS-GIS Centre |
| Road_cross_dist | Distance to nearest river-road crossing (km) | Own creation |
| **BFL** | Barrier Free Length, length of river between consecutive culverts (km) | Own creation |
| **BFL_share** | Proportion of total river length free of culverts | Own creation |
| Bioclimatic predictors |  |  |
| **Ap** | Presence of *Aplochiton* (y/n); coded no if unknown | Table S2 |
| **EW** | East or West Island | Own creation |
| **Slope** | Mean slope of grid cell | USGS |
| **Alt** | Mean altitude of each grid (m) | USGS |
| River_dens | River network in the basin (km) | Own creation, FIG IMS-GIS Centre |
| **Flow_accum** | Mean flow accumulation scaled by max flow accumulation in basin | Own creation, FIG IMS-GIS Centre |
| **Min_winter_temp** | Minimum winter temperature (°C) | FIG IMS-GIS Centre |
| **Rain** | Annual precipitation (mm) | FIG IMS-GIS Centre |
| **LC** | Land cover/substrate type | FIG IMS-GIS Centre, DPLUS065 Project |

## Notes

### Competing Interest Statement

The authors have declared no competing interest.

